# Uncovering the Diversity of Pathogenic Free-Living Amoebae in Freshwater Environments of Ghana: A Combined Culture Enrichment and Metabarcoding Approach

**DOI:** 10.1101/2025.05.17.654667

**Authors:** Yonas I. Tekle, Kwaku Oti Acheampong, Richard Kwame Adu, Kwaku Brako Dakwa

## Abstract

Free-living amoebae (FLA) such as *Naegleria* and *Acanthamoeba* are opportunistic pathogens increasingly linked to fatal and severe human infections, particularly in settings with limited hygiene, water sanitation and diagnostic infrastructure. In this pilot study, we explored the diversity of potentially pathogenic FLA in sectors of the Kakum River Basin, near Cape Coast, Ghana using a combined approach of filtered, pelleted, and culture-enriched metabarcoding. Our results revealed a wide range of FLA from Amoebozoa and Heterolobosea clades, including several of clinical relevance, such as *Acanthamoeba, Vermamoeba. Balamuthia* and *Paravahlkampfia*. Importantly, various *Naegleria* species were also detected and morphologically confirmed, raising public health concerns given the favorable environmental conditions for *Naegleria fowleri* proliferation in the sampling sites. The use of culture-enriched metabarcoding was particularly valuable in recovering organisms that may encyst and be missed by direct methods. This study underscores the importance of integrative and sensitive molecular approaches for detecting neglected pathogens in vulnerable communities. Our findings provide a foundation for larger epidemiological studies that incorporate clinical data and support the development of cost-effective monitoring tools for FLA-associated public health threats in sub-Saharan Africa.

---

Free-living amoebae (FLA) are ubiquitous protists (unicellular eukaryotes) found in diverse aquatic environments, such as rivers, lakes, and thermal springs, as well as in soil and dust (Page 1988; Schuster and Visvesvara 2004; Marciano-Cabral 2009). While most FLA are harmless, certain genera, including the Heterolobosean *Naegleria* and the amoebozoans *Acanthamoeba, Balamuthia, Vermamoeba* and *Sappinia*, can cause severe and often fatal infections such as Primary Amebic Meningoencephalitis (PAM) in humans (Gelman, et al. 2001; Visvesvara, et al. 2007; De Jonckheere 2012; Siddiqui and Khan 2014). PAM typically results from exposure to contaminated freshwater, where the amoeba enters the nasal cavity, migrates to the brain, and induces severe inflammation, leading to brain swelling and death within days (Yoder, et al. 2010). While PAM is rare, its high fatality rate (>97%) and the challenges involved in diagnosing and treating it make it a significant public health concern (Capewell, et al. 2015).

Cases of PAM have been reported in various African countries (Milanez, et al. 2023), including Nigeria, Zambia, and South Africa, often associated with swimming in warm freshwater, ritual ablution practices, or dust inhalation during the Harmattan season (Bhagwandeen, et al. 1975; Lawande, et al. 1979; Schoeman, et al. 1993; Chomba, et al. 2017). Despite these reports, no clinical cases of PAM or fatal FLA infections have been documented in Ghana to date, although the environmental conditions in the region may support the presence of pathogenic FLA. Given the increasing reports of *Naegleria* spp. and other FLA in neighboring regions, there is a pressing need to assess the potential risk posed by these organisms in Ghanaian water sources. A previous report in the Ashanti Region (Ghana) reiterates the potential existence of FLA in Ghanaian waters (De Jonckheere, et al. 2012).

A global systematic review of water sources across 35 countries revealed a pooled prevalence of 26.42% for *Naegleria* spp., with the pathogenic species *N. fowleri* detected in 23.27% of positive samples (Saberi, et al. 2020). These findings highlight both the widespread environmental distribution of pathogenic free-living amoebae and the urgent need for enhanced surveillance measures, particularly in regions with limited diagnostic capacity, high recreational water use, and warming aquatic environments that may favor pathogen proliferation (Cope and Ali 2016; Siddiqui, et al. 2016). Further, as wildlife biodiversity within the Kakum Conservation Area and its environs are sustained by the waters in the Kakum River Basin (KRB), the possibility of pathogenic FLA being caught up in the web of zoonosis cannot be overemphasized.

This study investigates the prevalence and diversity of pathogenic FLA in river water associated with communities in Twifo Hemang Lower Denkyira District, near Cape Coast in the Central Region, Ghana. Using a combined approach of culture enrichment and metabarcoding of environmental DNA (eDNA), we aimed to comprehensively assess the diversity of eukaryotic FLA present in these localities. Culture enrichment allows for the isolation of viable amoebae, while metabarcoding facilitates the detection of a broader range of FLA species, including those that may be missed by traditional methods. Through this integrated approach, we identify potential pathogenic species belonging to Heterolobosea (e.g., *Naegleria* spp.) and amoebozoans (e.g., *Acanthamoeba, Balamuthia* and *Vermamoeba*), which have been linked to human infections globally (Visvesvara, et al. 2007). Our findings reveal the presence of multiple FLA species implicated in human infections, as identified by the World Health Organization (WHO), in Ghanaian water sources.

This study represents a first thorough molecular-based environmental and biodiversity investigation of human-pathogenic FLA in Ghana. It provides valuable data on the distribution of these pathogens, which is crucial for assessing public health risks, especially in light of the increasing environmental changes and water pollution in the region. The results of this study will inform future surveillance efforts and guide the development of preventive strategies to mitigate the risks of waterborne FLA infections in Ghana.

## MATERIALS AND METHODS

### Water Sample Collection, Culture Enrichment, and Light Microscopy

Water samples were collected from two sites close to Cape Coast (in the Twifo Hemang Lower Denkyira District): Budamu Stream (5°16’27.348” N, 1°18’32.238” W) and Suruwi River (5°21’0.145”N and 1°23’0.583”W) during July and December of 2024. A total of five liters from each site were processed during each collection period for both metabarcoding DNA isolation and culture enrichment.

Three types of samples were collected for metabarcoding analysis: (1) direct-filtered environmental water, (2) detritus settled at the bottom of the water samples (hereafter referred to as “pellet”), and (3) culture-enriched samples. The culture-enriched samples were generated from multiple rounds of laboratory growth, as described below. Culture enrichment enables the proliferation of diverse microbes, including those underrepresented or in dormant cyst stages at the time of initial collection.

For the direct environmental sampling, water from each site was filtered using a 60 ml syringe and a handheld 47 mm diameter filter holder (Cobetter Polypropylene Filter Holder, 47FH-S) equipped with Whatman filters (Cytiva Whatman™ Cellulose Nitrate Membrane Filters) of varying pore sizes: 0.8 µm, 0.45 µm, and 0.2 µm. Filtering was performed sequentially from larger to smaller pore sizes to maximize the capture of both large and small microbes, as well as environmental DNA (eDNA). The remaining pellets from the water samples were processed further for eDNA extraction and live culture establishment. Filters and pellets were preserved in 2 ml of DNA lysis buffer for subsequent DNA extraction and analysis.

For the culture-enriched samples, a small portion of the pellet from each site was transferred into plastic Petri dishes. Live cultures were established by adding bottled natural spring water (Deer Park®, Nestlé Corp., Glendale, CA) along with autoclaved rice grains as a bacterial food source. After several rounds of subculturing, various microbial eukaryotes began to grow and were monitored. Samples from dishes showing robust microbial growth, including suspected free-living amoebae (FLA), were selected for metabarcoding analysis. Culture contents were scraped into 15 ml tubes and stored at −80°C. Over a period of 3–6 months, samples from each site were pooled and pelleted via centrifugation at 4000 rpm for 4 minutes. These pellets were then used for DNA extraction, as described below.

Monoclonal cultures of target species showing favorable growth were established on ATCC medium agar 997, a medium designed for freshwater amoebae. Morphological observations of isolates grown in plastic Petri dish cultures (ATCC medium agar 997) were made using an ECLIPSE Ti2 microscope (Nikon Corporation, Japan). Still images and videos were captured using the NIS-Elements software (Nikon Corporation) under phase contrast settings.

Measurements were conducted using the NIS software.

### DNA Extraction, PCR Amplification, and Sequencing

DNA extraction from preserved filters and pellets, originating from both natural and culture-enriched samples, was carried out using the Quick-DNA™ MicroPrep DNA Extraction Kit (Zymo Research, Irvine, California; Cat. No. D3020), following the manufacturer’s protocol. For pelleted samples, lysis buffer was added and thoroughly mixed by vortexing several times. The mixture was then centrifuged at 4000 rpm for 3 minutes, and the resulting clear supernatant was used for DNA extraction using the same kit. Total genomic DNA from clean monoclonal isolates, grown either in Petri dish cultures or on ATCC medium agar 997 and collected via pelleting, was also extracted using the same extraction protocol.

The full-length SSU-rDNA (18S) gene was amplified using primers described by Medlin, et al. (1988). For the metabarcoding study, primers targeting the V3–V4 region (KIM_18S_V4_forward : CCGCGGTAATTCCAGCTC and Hadziavdic_18S_V4_revsrse: CCCGTGTTGAGTCAAATTAAGC) of the 18S gene were used (Nickrent and Sargent 1991; Hadziavdic, et al. 2014). Phusion DNA polymerase, a high-fidelity proofreading enzyme, was used for PCR amplification under conditions described in (Tekle, et al. 2013). To capture a broad diversity of amplicons from various eukaryotic groups, independent PCR reactions were performed with annealing temperatures ranging from 50°C to 60°C. PCR amplicons using the were purified using the QIAquick PCR Purification Kit (Qiagen, Cat. No. 28106) and pooled to ensure sufficient DNA concentration for Oxford Nanopore sequencing. DNA concentrations were quantified using a Qubit assay with the dsDNA Broad Range Kit (Life Technologies, Carlsbad, CA, USA).

Library preparation was performed using the Oxford Nanopore Technologies Rapid Barcoding Kit (SQK-RBK114.24), according to the manufacturer’s instructions. Prepared libraries were loaded onto an R10.4.1 flow cell and sequenced using a MinION device. Raw reads were processed and prepared for downstream analysis as described in (Tekle, et al. 2025).

Additionally, amplicons derived from monoclonal cultures were sent to Azenta Life Sciences (Burlington, MA, USA) for sequencing using their PCR-EZ platform. Consensus sequences from these monoclonal samples were generated using Azenta’s proprietary bioinformatics pipeline.

### Metabarcoding phylogenetic and pairwise analyses

To assess eukaryotic diversity, Operational Taxonomic Units (OTUs), in environmental samples a custom nucleotide BLAST database was built, from the SILVA 138.2 SSU rDNA database (Quast, et al. 2012), using the makeblastdb utility from the BLAST+ suite (v2.6.0). All sequencing reads were run locally against this database using Blastn with the parameters - max_target_seqs 1 and an e-value threshold of 1e-10, to only retain the top and highest-confidence hit per read. Taxonomic ranks were extracted from the relevant columns of the combined BLAST output. Sample metadata, including the sampling sites (Budamu Stream or Suruwi River) and sample types (Filter, Pellet, or Enriched), were parsed from sequence identifiers embedded in the sample names. Data processing and visualization were conducted using the dplyr and ggplot2, packages in R version 4.4.2.

The molecular phylogeny of the 18S rDNA gene from *Naegleria* species was reconstructed using IQ-TREE (Nguyen, et al. 2015; Kalyaanamoorthy, et al. 2017; Hoang, et al. 2018), based on alignments generated in AliView (Larsson 2014). The best-fit evolutionary model was selected automatically using the -m AUTO option, and node support was assessed with 1,000 ultrafast bootstrap replicates (-B 1000). The alignment used for phylogenetic tree reconstruction included 3,383 nucleotide sites comprising 43 ingroup and one outgroup taxon. A similar alignment, trimmed for maximum overlap (3,280 nucleotide sites), was used to compute pairwise genetic distances in MEGA (Kumar, et al. 2001) using the Kimura 2-Parameter (K2P) model.

## RESULTS

### Sampling Sites and Overall Eukaryotic Microbial Diversity

The two sampling sites, Budamu Stream and Suruwi River, are part of a broader research project investigating human settlements along the Kakum River basin in the Central Region of Ghana, West Africa. The Suruwi River provides water for domestic and agricultural use to residents of Frami Township (Twifo Hemang Lower Denkyira District), whilst the Budamu Stream, located upstream, serves the Abrafo-Odumasi communities (Twifo Hemang Lower Denkyira District). Both water bodies are frequently used for bathing, drinking, and washing by locals and they are tributaries of the Kakum River that passes through the Kakum Conservation Area near Cape Coast in the central region. Ongoing assessments are investigating the trends of pollution levels resulting from anthropogenic activities, especially agricultural runoffs, fertilizer usage along the KRB and their implications on microbial diversity as well as distribution.

In this study, we report on the overall microbial diversity and the prevalence of free-living amoebae (FLA) in these aquatic environments. Microbial community profiles were generated using metabarcoding of the 18S gene (V3/V4 variable regions). Both sites exhibited comparable microbial diversity, with dominant taxonomic groups belonging to the Alveolata and Obazoa clades (Figure 1). Budamu Stream showed a higher relative abundance of Obazoa and a lower abundance of Alveolata compared to Suruwi River (Figure 1). Within the Alveolata, ciliates accounted for approximately 99% of the observed diversity, whereas the Obazoa group was dominated by rotifers and gastrotrichs. Microscopic examination of both fresh and enriched cultures confirmed the presence of these taxa. Other groups including Amoebozoa, Chloroplastida, Cryptomonadales, Rhizaria, Stramenopiles, and Discicristata were present at similar levels across both sites (Figure 1). A more comprehensive diversity analysis of these sites will be presented in future studies.

**Figure 1.**
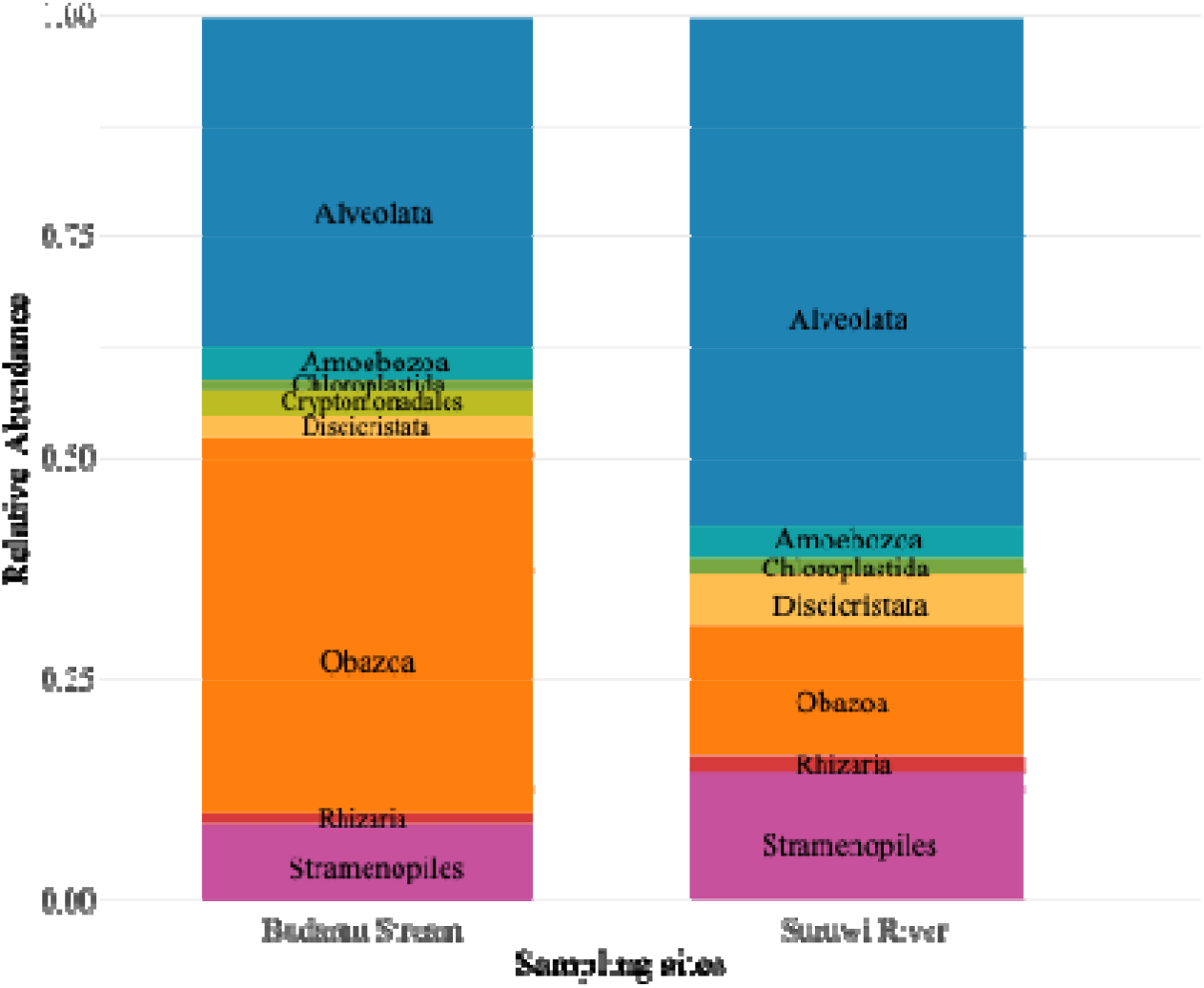
Relative abundance of top eukaryotic phyla across sampling sites.

### Discovery and Diversity of Free-Living Amoebae (FLA) Across Sites and Sample Types

A more focused metabarcoding analysis of FLA diversity using different sample types (filtered, pelleted, and culture-enriched) revealed notable differences in taxonomic composition between the two sites (Figures 2 and 3). Known FLA lineages from Heterolobosea (Discoba) and Amoebozoa were recovered from both Budamu Stream and Suruwi River, although the taxonomic breadth varied across sample sites and types. Overall, Budamu Stream exhibited greater FLA diversity than Suruwi River (Figures 2–5). Culture-enriched samples consistently revealed higher taxonomic diversity than freshly filtered or pelleted samples (Figures 2 and 3).

**Figure 2.**
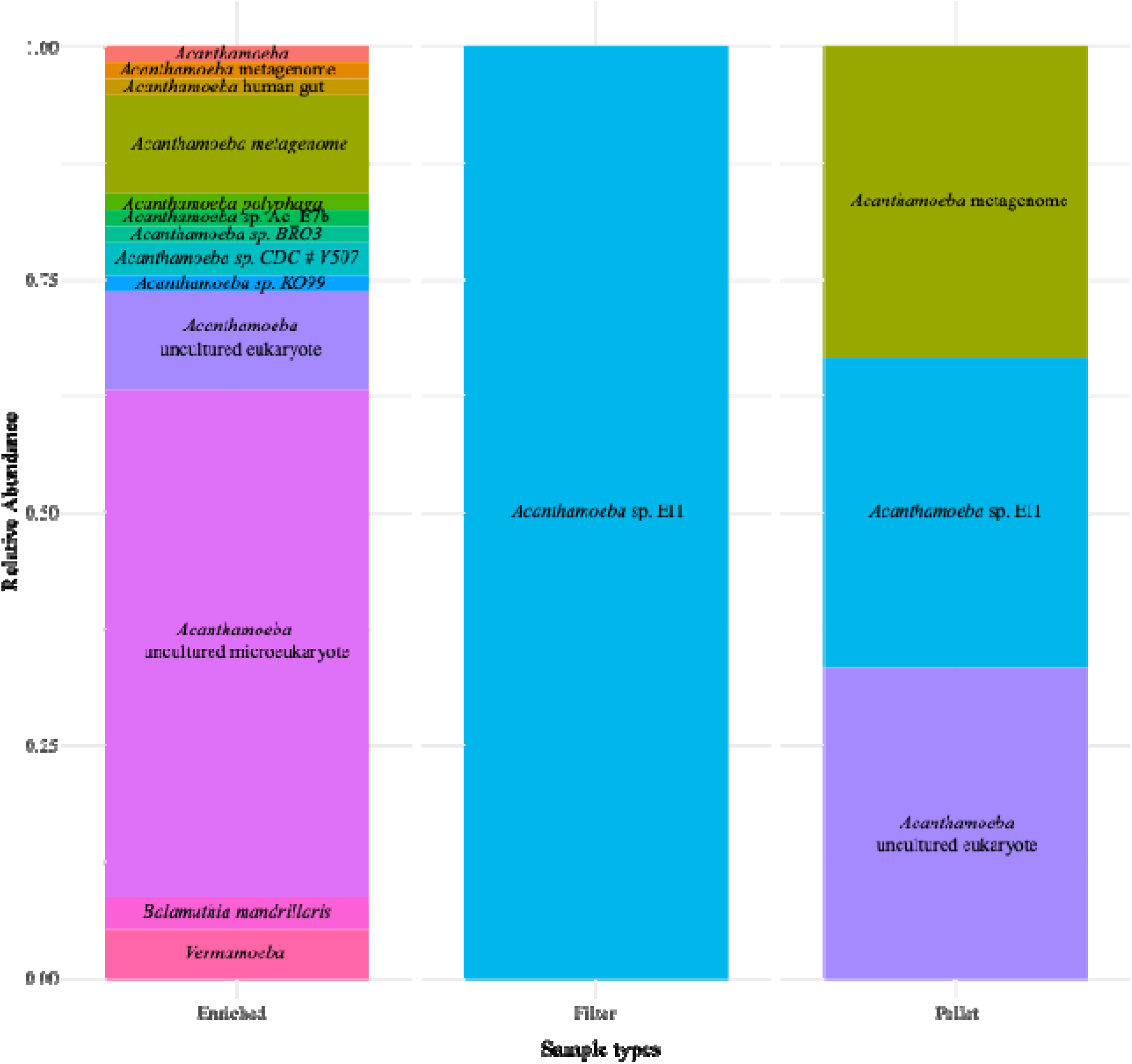
Relative abundance of free-living amoebae (FLA) belonging to Amoebozoa in Budamu Stream, including cultured-enriched, filter, and pellet samples. Note the high diversity in the cultured-enriched sample.

**Figure 3.**
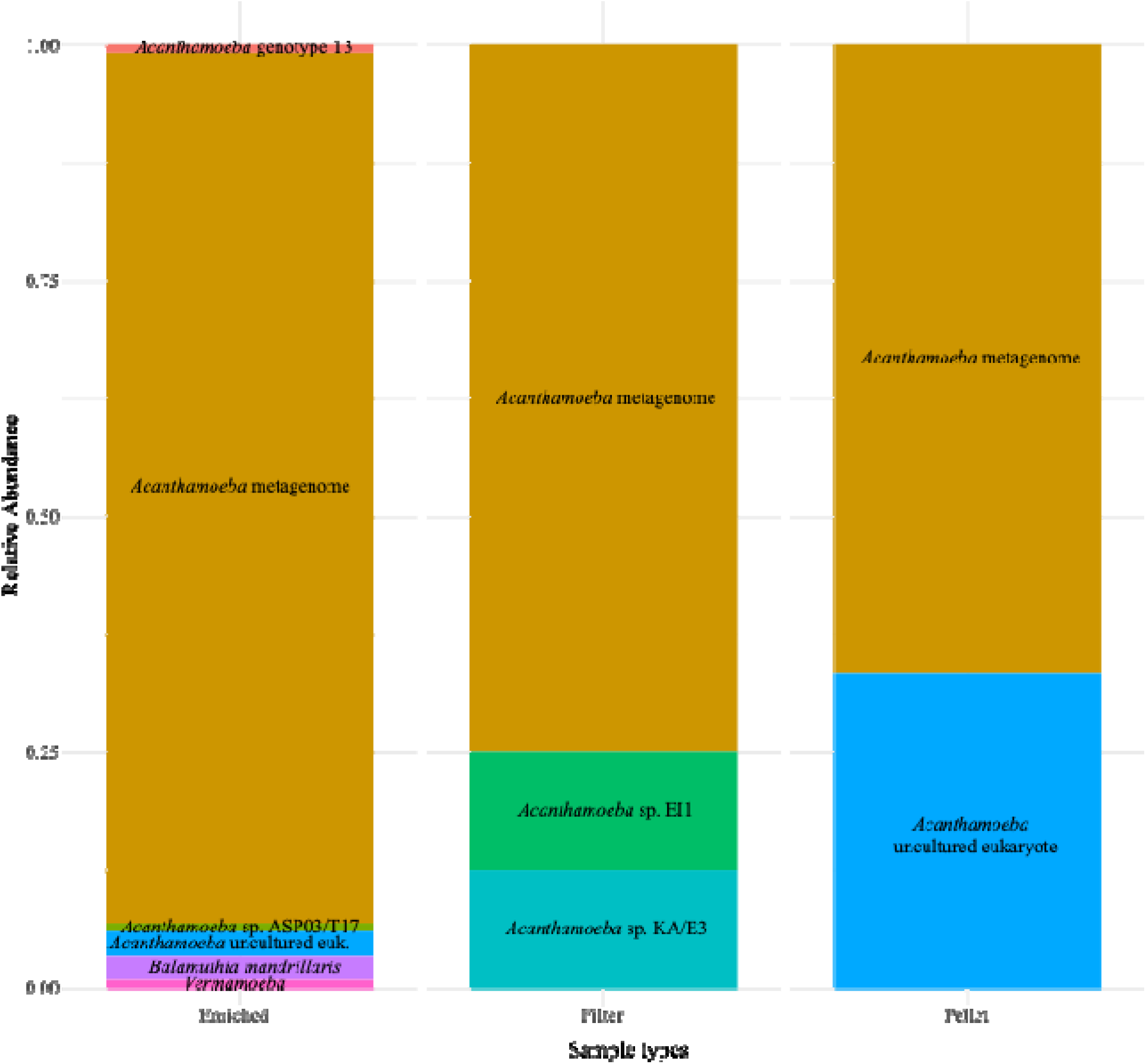
Relative abundance of free-living amoebae (FLA) belonging to Amoebozoa in Suruwi River, including cultured-enriched, filter, and pellet samples.

Almost all the FLA genera (*Acanthamoeba, Balamuthia*, and *Vermamoeba*) except *Sappina* from Amoebozoa implicated in human infections, were detected. Only *Acanthamoeba* lineages were found in the freshly processed samples (filtered and pelleted), whereas all three amoebae genera were recovered from culture-enriched samples at both sites (Figures 2 and 3). Budamu samples generally exhibited more diverse *Acanthamoeba* strains and a slightly higher overall abundance of FLA (Figure 2).

A similar biodiversity pattern was observed among Discoba (Excavata) FLA. Culture-enriched samples revealed most of the relevant FLA, while filtered and pelleted samples showed limited diversity (data not shown). In total, 13 genera belonging to Euglenozoa and Heterolobosea were identified in culture-enriched samples (Figures 4 and 5). Among them were human-pathogenic genera such as *Naegleria* and *Paravahlkampfia* (Figure 4), as well as potential animal or fish pathogens like *Ichthyobodo*. Taxonomic differences between the two sites were evident: all Discoba taxa from Budamu Stream belonged to Heterolobosea, except for *Petalomonas*, an Euglenozoan (Figure 4). Conversely, the Suruwi River samples contained predominantly non-pathogenic Euglenozoans, with two exceptions: *Ichthyobodo*, a known fish parasite, and *Allovahlkampfia*, a free-living heterolobosean (Figure 5). Additional unclassified environmental Discoba species were also identified across both sites (Figures 4 and 5).

**Figure 4.**
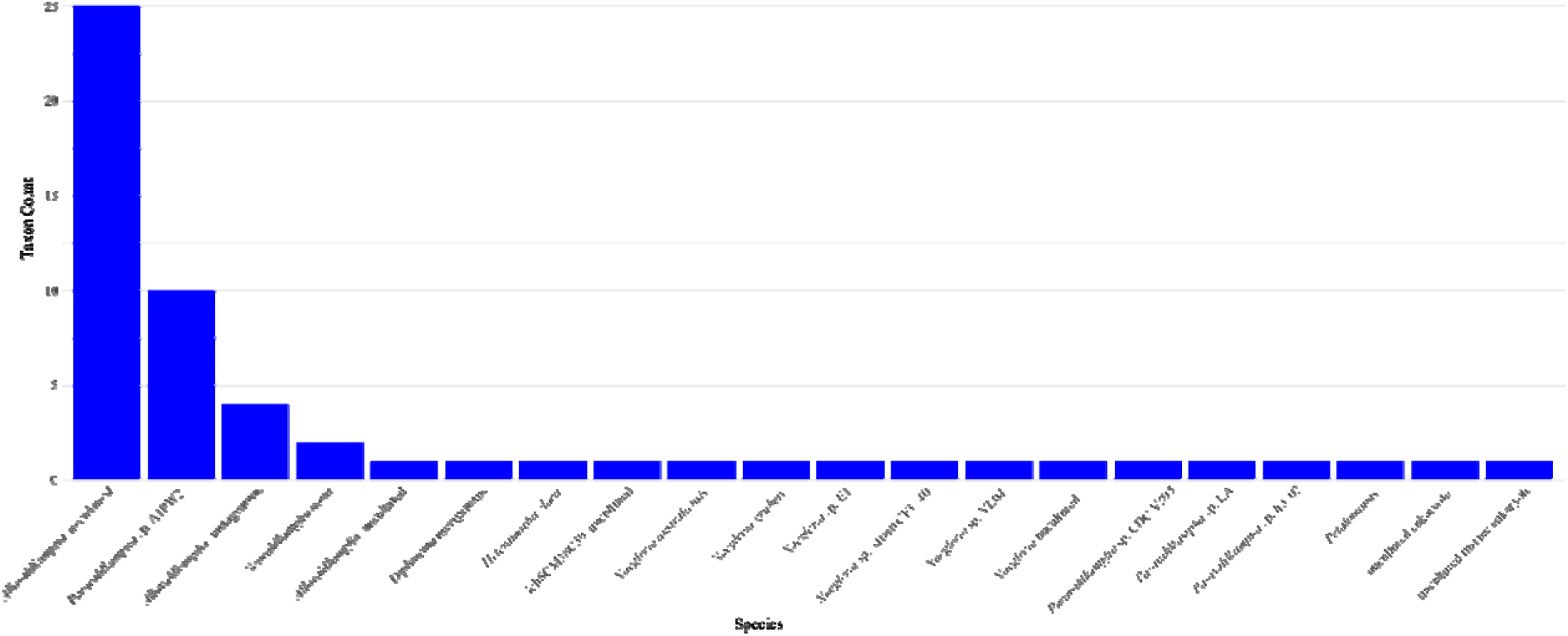
Relative abundance of free-living amoebae (FLA) belonging to Discoba in Budamu Stream, based on cultured-enriched samples. Note the diverse and high recovery of *Naegleria* spp.

**Figure 5.**
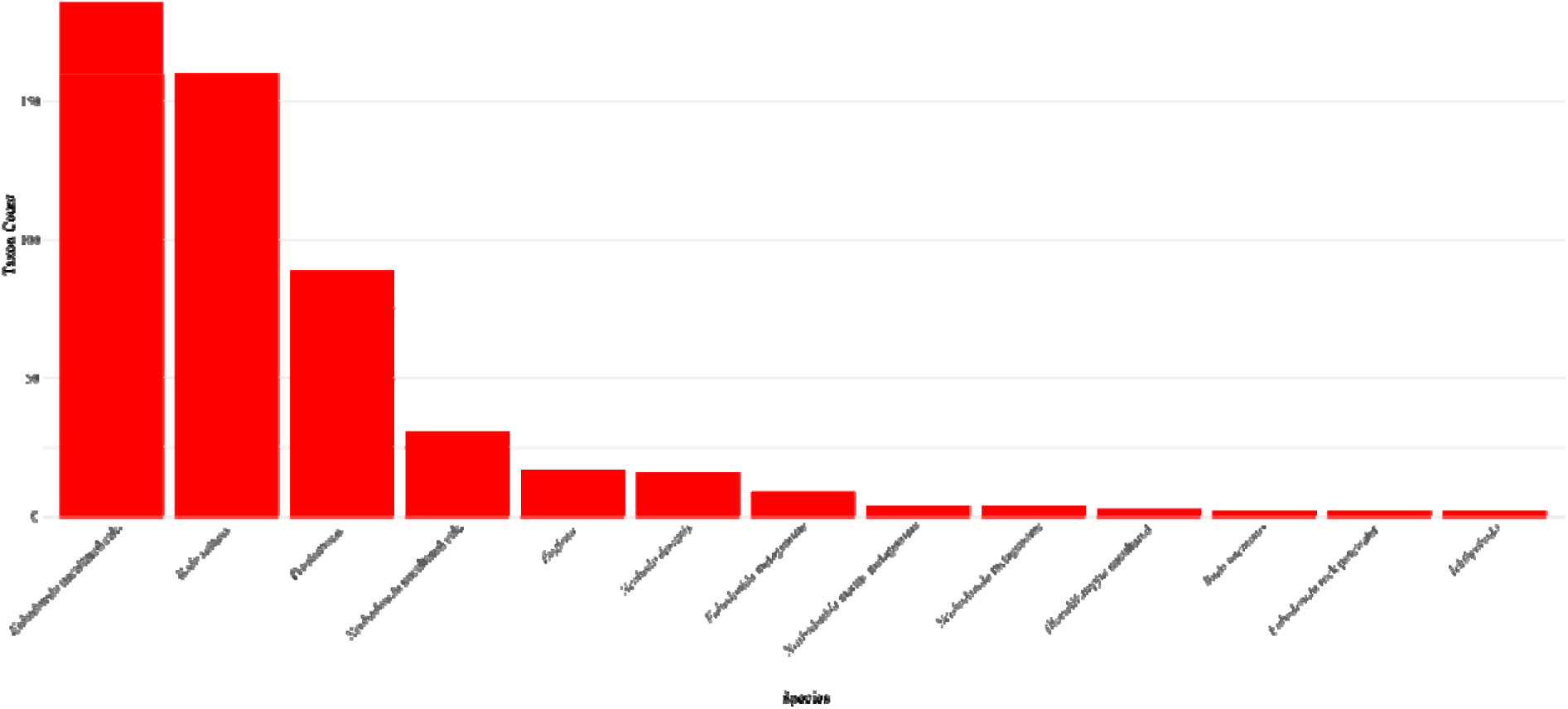
Relative abundance of Discoba in Suruwi River.

### Culture-Based Diversity Assessment and Isolation of *Naegleria* sp

To complement the metabarcoding approach, we performed culture-based assessments from environmental samples. These confirmed the presence of diverse ciliates, rotifers, and gastrotrichs detected in the sequencing data. In addition, culturing revealed FLAs that were not captured through direct metabarcoding, supporting the observation that culture-enriched samples provide deeper insights into microbial diversity.

Notably, a relative of the pathogenic species *Naegleria fowleri* was successfully isolated and monoclonally cultured. The isolate, designated *Naegleria* sp. YT4-GH, displayed classic heterolobosean eruptive pseudopodal locomotion. Cells averaged 41 µm in length and 23 µm in width (Figure 6). During active movement, they exhibited a characteristic “worm-like” (finger-shaped) morphology, with occasional triangular, round, or branching forms (Figure 6). Trailing filaments at the posterior end of the cells were observed during locomotion (Figure 6D, E). No flagellated stage was identified. The cysts, averaging 12 µm in diameter, showed a dark central body surrounded by a bright outer halo (Figure 6I).

**Figure 6.**
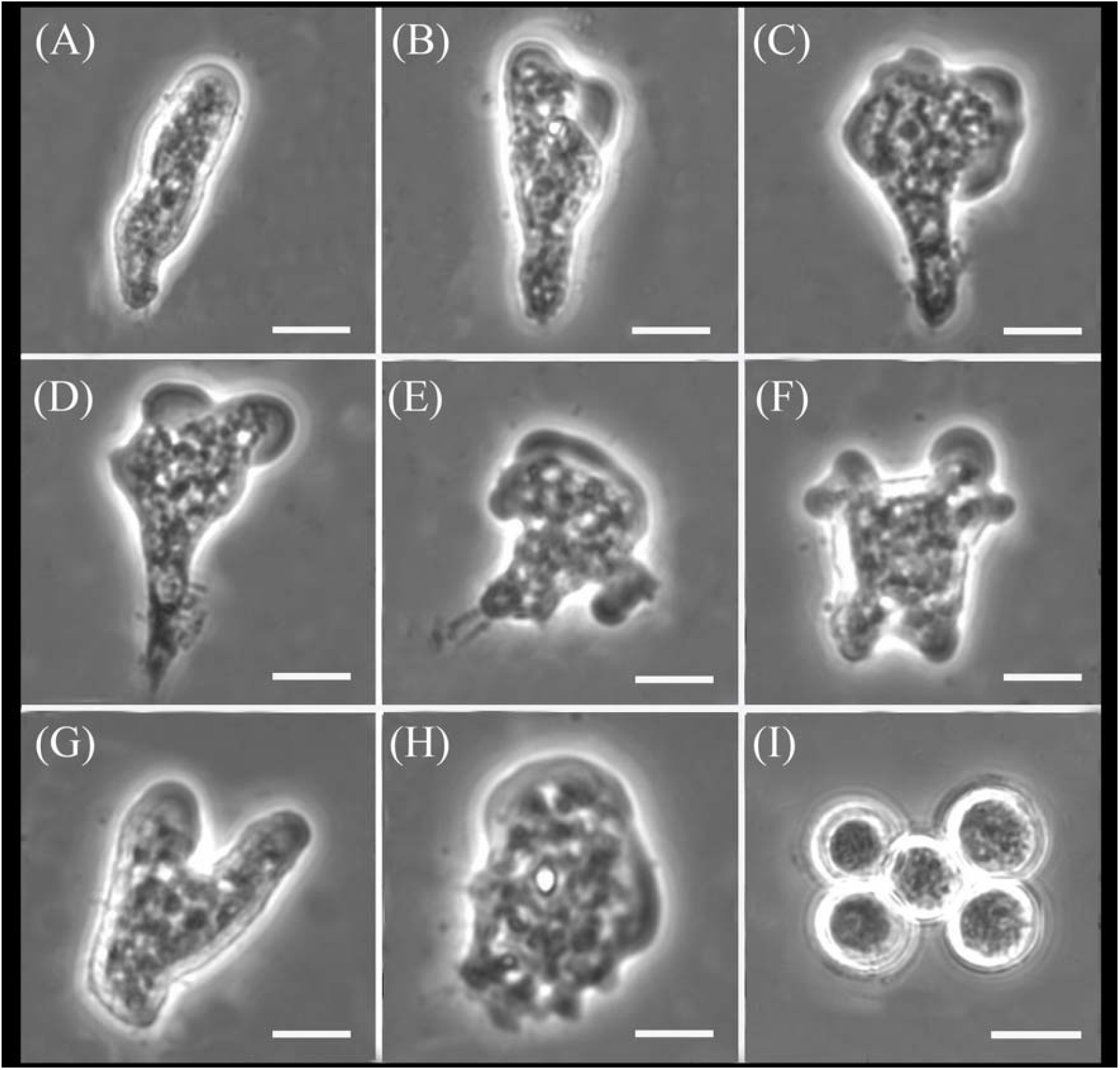
Phase contrast micrographs of *Naegleria* sp. YT4-GH showing different locomotive forms (A–H) and cyst (I). All scale bars represent 10 µm.

Molecular phylogenetic analysis of the 18S gene placed the YT4-GH isolate within the *Naegleria clarki* clade, with strong bootstrap support (Figure S1). Pairwise divergence between the YT4-GH isolate and other *N. clarki* strains ranged from 0.06% to 0.43% (Table S1), confirming its close genetic affiliation with this species. The genus *Naegleria* is generally characterized by low divergence in the 18S rDNA gene; for instance, *Naegleria fowleri*, a more distantly related species, differs from YT4-GH by only 6.04% (Table S1). Based on both morphological and molecular data, the YT4-GH isolate is most likely *N. clarki*, although further confirmation using faster-evolving genetic markers such as the internal transcribed spacer (ITS) region is recommended. The successful isolation of the *Naegleria* YT4-GH strain complements our metabarcoding findings and reinforces the utility of culture-enriched sampling as a reliable method for comprehensive assessment of FLA diversity.

## DISCUSSION

### Metabarcoding detection of Neglected but Dangerous Pathogens in Resource-Limited Settings

Free-living amoebae (FLA) such as *Naegleria* and *Acanthamoeba* are increasingly recognized as opportunistic human pathogens, capable of causing severe and often fatal infections. In resource-limited settings, particularly in parts of sub-Saharan Africa where recreational and domestic use of freshwater bodies is common and poorly regulated, the public health threat posed by these organisms remains largely overlooked (Chomba, et al. 2017; Milanez, et al. 2023). This study represents one of the few investigations in West Africa that uses high-throughput sequencing tools to examine the environmental diversity of FLA in water bodies actively used by local communities.

Historically, studies on free-living amoebae (FLA) in Africa have been limited, often relying on traditional microscopy, culture-based techniques, and a few PCR-based methods (reviewed in (Milanez, et al. 2023). Most reports have come from North African countries particularly Egypt, with additional cases reported in Algeria and Tunisia (Dendana, et al. 2013; AI-Herrawy 2014; Al-Herrawy and Gad 2015; Chomba, et al. 2017). Sporadic reports from other African countries have identified FLA in both clinical and environmental samples (Lawande, et al. 1979; Lastovica 1980; Eddyani, et al. 2008; van der Beek, et al. 2015; Sente, et al. 2016; Chomba, et al. 2017; Saberi, et al. 2020). However, data from most West African countries and especially Ghana remain scarce. Existing studies have typically focused on isolated clinical cases or limited geographic regions. This study broadens the scope by applying a metabarcoding-based approach, enabling more comprehensive detection of FLA diversity directly from environmental DNA (eDNA).

The detection of multiple potentially pathogenic amoebae from samples collected in close proximity to human activity zones underscores a pressing need for broader surveillance. The application of eDNA techniques in this context allows for the detection of pathogens without the need for invasive sampling or laboratory-dependent diagnostics (Sengupta, et al. 2019). This is especially critical in regions with constrained healthcare infrastructure. The findings from this pilot study therefore serve as a foundation for future work that will incorporate clinical surveillance, seasonal sampling, and intervention strategies. A more extensive study, with integration of both environmental and clinical datasets, could enable the development of cost-effective tools for monitoring and mitigating the risks of FLA-related infections.

### Advantages of Multi-Approach Metabarcoding

This study also provides a methodological contribution by evaluating and comparing three distinct metabarcoding workflows: direct filtered water eDNA, pelleted eDNA, and culture-enriched eDNA. Metabarcoding studies, while powerful, are known to be influenced by sample processing methods, PCR biases, and sequencing depth, which can lead to under-or overestimation of certain biodiversity (Elbrecht and Leese 2015; Alberdi, et al. 2018; Catlett, et al. 2020). By employing multiple parallel approaches, we were able to capture a wider range of FLA taxa than would have been possible with a single technique.

Most notably, the use of culture-enriched metabarcoding played a critical role in recovering taxa that might otherwise be missed in direct eDNA workflows. FLA, especially during periods of environmental stress, often encyst, a process that can make them resistant to lysis and thereby invisible to standard DNA extraction methods. Our culture enrichment step allowed these cysts to excyst and proliferate under favorable laboratory conditions before DNA extraction, thereby improving detection rates. This approach not only enhanced sensitivity but also allowed for the growth and isolation of viable FLA, which is essential for downstream phenotypic and pathogenicity assays.

The clear complementarity between the culture-enriched and direct methods reinforces the importance of methodological diversity in environmental pathogen studies. Future environmental surveillance programs, particularly those in resource-limited regions, should consider integrating culture-enriched workflows as a standard component when assessing the risk posed by FLA and similar organisms.

### Isolation of *Naegleria* and observation of Pathogenic Amoebozoans

The successful isolation of *Naegleria* from our environmental samples is an important and concerning finding. Although the exact species isolated in this study may not be the cause of amoebic meningoencephalitis (PAM, *Naegleria fowleri*), the isolation and discovery of multiple species of *Naegleria* spp. in the samples particularly Budamu Stream highlights the suitability of the studied water bodies to support *Naegleria* populations. Given that *Naegleria fowleri* shares similar environmental niches particularly warm, stagnant, and untreated freshwater sources, the detection of any *Naegleria* species signals a potential risk (De Jonckheere 2012). Water temperatures recorded at the time of sampling were well within the optimal range for the growth and proliferation of *Naegleria fowleri* (25°C to 46°C), suggesting that conditions are favorable for the emergence or persistence of the pathogenic strain. Similarly, the detection of *Paravahlkampfia* sp. CDC #V595 further corroborates the existence of FLA that cause infection in Humans. *Paravahlkampfia* sp. CDC #V595 was isolated from the cerebrospinal fluid of a patient and causes PAM similar to *Naegleria fowleri* (Visvesvara, et al. 2009).

Moreover, our study identified in culture several amoebozoan taxa known or suspected to be pathogenic, including *Acanthamoeba*, *Balamuthia*, and *Vermamoeba* (data not shown). The observation of these potential pathogens in culture supports and reinforces their detection through metabarcoding analysis. These free-living amoebozoans (FLA) are associated with a range of severe to fatal infections, including *Balamuthia amoebic encephalitis* (BAE), *granulomatous amoebic encephalitis* (GAE), *Acanthamoeba keratitis* (AK), and cutaneous acanthamoebiasis (Naginton, et al. 1974; Martinez, et al. 1994; Visvesvara, et al. 2007; Bravo and Gotuzzo 2016). Additionally, these FLA are known to harbor intracellular pathogens such as *Legionella* and *Mycobacterium*, effectively acting as “Trojan horses” that facilitate the survival, spread, and even evolution of emerging microbial threats (Molmeret, et al. 2005). The presence of these organisms in water sources used for recreation, household washing, or even drinking underscores the urgent need for routine environmental monitoring and public health education to prevent potentially life-threatening infections.

## Conclusion

This study is among the first in Ghana and one of the few in sub-Saharan Africa to use high-throughput, multi-modal metabarcoding to examine free-living amoebae in freshwater environments frequented by the public. The combination of pelleted, filtered, and culture-enriched DNA workflows provided a robust assessment of FLA diversity, uncovering taxa of clinical significance and demonstrating the superiority of multi-pronged approaches. The detection of *Naegleria* spp., as well as other CDC-recognized pathogenic isolates (*Acanthamoeba* and *Paravahlkampfia*), illustrates the real and present public health risks associated with untreated water exposure in these areas.

These findings call for expanded surveillance, including temporal sampling and clinical correlations, to accurately assess risk and inform public health strategies. Moving forward, our results provide a compelling basis for the development of low-cost diagnostic and monitoring tools that can be applied in other endemic or under-studied regions to prevent avoidable morbidity and mortality associated with FLA infections.

## ACKNOWLEDGEMENTS

This work was supported by the National Science Foundation EiR award (2401946) and the Simons Fellow Award (SFA-23-5) to Y.I.T. Additional travel support was provided by the NSM Faculty Seed Award and the Gordon-Zeto Center for Global Education at Spelman College. We thank Joshua Kobby Erskine for his assistance with sample collection and processing, and Saron Ghebezadik and Priyal Patel for their help with data analysis. We are also grateful to Dr. Rofela Combey for facilitating and organizing the field trips and our visit to the University of Cape Coast.

## Supplementary Figure

**Figure S1.**
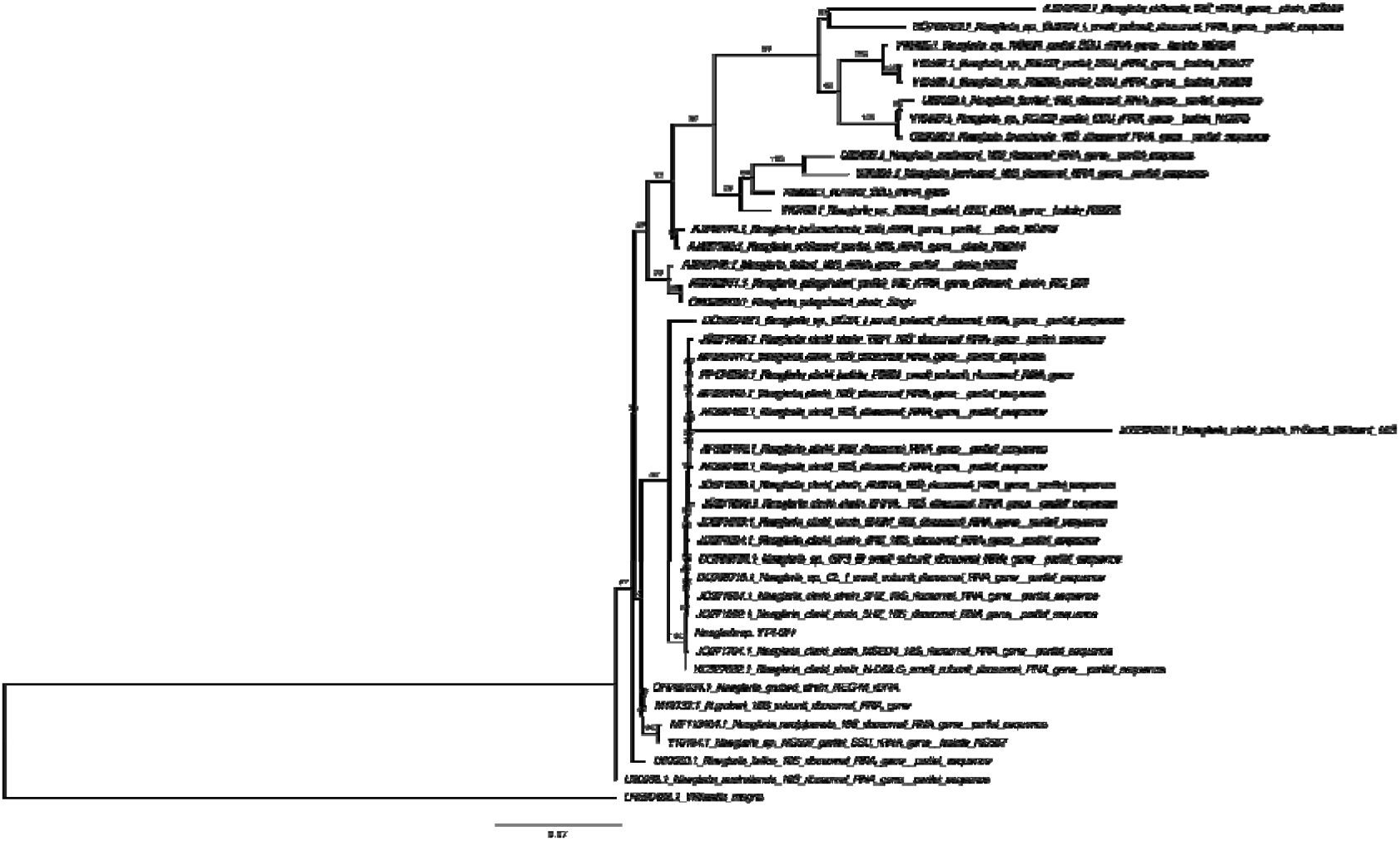
IQ-TREE maximum likelihood phylogeny of the 18S rDNA gene based on 43 ingroup taxa of the genus *Naegleria*, showing the placement of *Naegleria* sp. YT4-GH. Numbers at nodes represent IQ-TREE bootstrap values. Branches are drawn to scale.

**Table S1.** Pairwise distances of 18S gene of our isolate Naegleria sp. YT4-GH and other Naegleria spp.

| Species Pair | Min Divergence | Max Divergence | Avg Divergence |
| --- | --- | --- | --- |
| M18732.1 *Naegleria gruberi* vs *Naegleria* sp. YT4-GH | 0.61% | 0.61% | 0.61% |
| Y10184.1 *Naegleria* sp. NG597 vs *Naegleria* sp. YT4-GH | 0.76% | 0.76% | 0.76% |
| U80060.1 *Naegleria italica* vs *Naegleria* sp. YT4-GH | 1.13% | 1.13% | 1.13% |
| U80058.1 *Naegleria australiensis* vs *Naegleria* sp. YT4-GH | 1.33% | 1.33% | 1.33% |
| AJ243440.1 *Naegleria fultoni* NG885 vs *Naegleria* sp. YT4-GH | 1.66% | 1.66% | 1.66% |
| AM062741.1 *Naegleria pringsheimi* NG007 vs *Naegleria* sp. YT4-GH | 2.21% | 2.21% | 2.21% |
| AJ237786.1 *Naegleria robinsoni* NG944 vs *Naegleria* sp. YT4-GH | 2.31% | 2.31% | 2.31% |
| AJ243441.1 *Naegleria indonesiensis* NG945 vs *Naegleria* sp. YT4-GH | 2.70% | 2.70% | 2.70% |
| X93224.1 *Naegleria minor* vs *Naegleria* sp. YT4-GH | 4.21% | 4.21% | 4.21% |
| U80061.1 *Naegleria jamiesoni* vs *Naegleria* sp. YT4-GH | 5.61% | 5.61% | 5.61% |
| U80062.1 *Naegleria lovaniensis* vs *Naegleria* sp. YT4-GH | 5.71% | 5.71% | 5.71% |
| U80057.1 *Naegleria andersoni* vs *Naegleria* sp. YT4-GH | 5.85% | 5.85% | 5.85% |
| U80059.1 *Naegleria fowleri* vs *Naegleria* sp. YT4-GH | 6.04% | 6.04% | 6.04% |
| Y10189.1 *Naegleria* sp. NG055 vs *Naegleria* sp. YT4-GH | 7.33% | 7.33% | 7.33% |

